# Emotion Representation and Neural Synchrony: Decoding Valence and Arousal with Wearable EEG

**DOI:** 10.64898/2026.06.03.730031

**Authors:** Iwon Yang, Chaery Park, Jongwan Kim

**Affiliations:** Pusan National University, Pusan-si, Republic of Korea; Jeonbuk National University, Jeonju-si, Jeonbuk State, Republic of Korea

**Keywords:** wearable EEG, core affect, valence, arousal, classification, multidimensional scaling, intersubject correlation

## Abstract

Emotions are dynamic experiences that unfold over time, yet most affective neuroscience studies have relied on static stimuli and laboratory-based EEG systems. This study examined whether emotional valence and arousal can be reliably decoded using a consumer-grade wearable EEG device in naturalistic contexts. Forty-three participants viewed video clips designed to elicit four core affect categories including high-arousal positive, low-arousal positive, high-arousal negative, and low-arousal negative, while EEG signals were continuously recorded. Multivariate analyses, including classification, multidimensional scaling (MDS), and intersubject correlation (ISC), were employed to assess affective representation and neural synchrony. Behavioral data demonstrated robust classification of both valence and arousal, whereas EEG data yielded consistent above-chance classification of valence but less stable decoding of arousal, particularly in within-participant analyses. MDS revealed that both behavioral and EEG responses were primarily organized along the valence dimension, with weaker separation along arousal. ISC analyses further indicated frequency- and region-specific neural synchrony, with stronger alignment in left and temporal electrodes, though overall ISC values were modest, likely reflecting the brief duration of stimuli. Taken together, these findings suggest that valence is more stably represented in both subjective and neural domains, whereas arousal may require time-resolved or longer-duration approaches for reliable decoding. This work demonstrates the feasibility and limitations of employing wearable EEG for theory-driven affective neuroscience, underscoring its potential for scalable and ecologically valid emotion research beyond laboratory settings.

## 1. Introduction

Emotions are temporally unfolding phenomena, often embedded within dynamic environments such as film, music, or social interaction (Gross & Levenson, 1995; Kreibig, 2010). Compared to static or decontextualized stimuli, these naturalistic contexts evoke affective experiences that evolve over time, involving fluctuations in valence and arousal. Understanding such dynamic responses is essential for capturing how emotions are processed and regulated in real-life situations. To address this, affective science has increasingly adopted naturalistic stimuli such as emotionally evocative video clips (Hasson et al., 2004). These types of stimuli offer immersive and context-rich affective content, making it possible to study emotion in settings that more closely resemble everyday life (Nastase et al., 2019; Golland et al., 2014). They also allow researchers to examine not only the presence of emotional states but also how these states unfold over time and synchronize across individuals. To investigate the neural underpinnings of emotional experiences, many studies have employed electroencephalography (EEG). EEG has been a prominent method for capturing the temporal dynamics of affective responses, providing millisecond-level resolution of neural activity. Numerous studies have utilized EEG in controlled laboratory settings. Laboratory-grade EEG systems have allowed researchers to examine affective processes with high temporal resolution, frontal alpha asymmetry, and other markers of emotional reactivity (Davidson, 1992; Coan & Allen, 2004). These methods have yielded important insights into how emotional valence and arousal are represented in the brain.

Despite these advantages, EEG research in controlled laboratory settings faces important limitations. Traditional EEG systems typically require costly equipment, extensive preparation, and strictly controlled environments, which restrict their ecological validity and accessibility in naturalistic contexts (Debener et al., 2012; Ratti et al., 2017).These systems often involve time-consuming setup procedures, such as scalp abrasion and gel application, which can cause discomfort and make them unsuitable for certain populations, including elderly or clinical groups with limited tolerance for long procedures (Ratti et al., 2017). Moreover, while traditional multi-lead systems are advantageous in terms of signal quality and analytical depth, their low portability and poor usability make them ill-suited for applications requiring rapid deployment, such as field studies or large-scale emotion tracking.

To overcome these limitations, recent advances in wearable EEG technology have enabled the collection of brain data in more flexible and naturalistic contexts. Consumer-grade devices (i.e. wearable devices) offer a lightweight, cost-effective alternative that can be used in ambulatory environments while still providing usable EEG signals for emotion research. Although these devices have fewer channels and lower signal resolution compared to lab-based systems, their accessibility and convenience make them a promising tool for real-world affective neuroscience. The Muse 2 (InteraXon Inc.) shows lower reliability compared to medical devices, but it is generally considered sufficient for use in nonclinical research settings (Ratti et al., 2017). It has also been successfully applied to quantify event-related brain potentials (Krigolson et al., 2017) and to detect emotional responses (Raheel et al., 2020). In line with these advantages, the dataset provided by Saganowski et al. (2022) demonstrated the feasibility of using consumer-grade wearable devices, such as Muse 2, to elicit and measure emotional responses in naturalistic settings. This dataset included multimodal physiological signals and self-reported affective ratings in response to short film clips designed to evoke discrete emotions. Building on this foundation, Yang et al. (2024) reanalyzed the dataset and examined affective responses in depth, confirming that EEG data can successfully distinguish between emotional states.

Multivariate analysis of EEG provides the sensitivity to detect subtle neural differences and, with high spatial and temporal precision, allows for the fine-grained identification and continuous tracking of specific stimulus features, thereby offering a powerful approach for analysis (Fahrenfort et al., 2017; Hascher et al., 2023). Classification analysis and multidimensional scaling (MDS) are particularly well-suited for affective EEG research, as they enable the identification of latent representational structures and predictive patterns across complex datasets. MDS allows for a spatial visualization of affective states in a low-dimensional space, while classification models assess the separability of emotion categories based on EEG features. Together, these methods facilitate a more nuanced understanding of how affect is encoded in the brain and can be used to validate theoretical emotion models empirically.

While the primary goal of this study was to examine whether affective states can be systematically represented and classified using EEG signals collected from a wearable device, we additionally employed intersubject correlation (ISC) analysis to explore the temporal synchronization of neural responses across participants. Given the high temporal resolution of EEG, this modality enables the investigation of time-locked neural dynamics that unfold during emotional video viewing (Nastase et al., 2019). Incorporating ISC as a secondary analysis thus allowed us to assess shared affective processing from a complementary, time-resolved perspective. This not only complements the classification and representational results, but also enables a cross-validation of individual and shared neural processes. Although ISC has been primarily applied to fMRI data, it can also be used across other imaging modalities such as EEG and fNIRS. AS a high temporal resolution modality, EEG is particularly well-suited for capturing stimulus-evoked dynamics in ISC frameworks (Nastase et al., 2019; Dmochowski et al., 2012). Neural activity unfolds at different temporal scales, and EEG enables the measurement of fine-grained, rapid fluctuations in response to dynamic affective stimuli. This temporal precision complements the slower hemodynamic signals measured by fMRI. However, as ISC is modulated by the narrative structure and temporal coherence of the stimulus (Dmochowski et al., 2012), EEG-based ISC values may vary considerably depending on experimental design and signal quality.

Building on previous findings, the present study aimed to determine whether affective representation and classification are feasible using a consumer-grade wearable EEG device (Muse 2). By organizing emotional stimuli along the valence-arousal space into four core affect categories (High-arousal Positive, Low-arousal Positive, High-arousal Negative, Low-arousal Negative), we aimed to reduce ambiguity and increase the interpretability and discriminability of affective EEG responses. This dimensional approach also aligns with previous affective neuroscience research emphasizing the continuous structure of affective space (Russell, 1980; Russell & Barrett, 1999). Moreover, we introduced ISC analysis to assess the degree of neural response consistency across participants. By quantifying EEG synchronization during emotional video viewing, we sought to investigate whether shared affective experiences elicit shared neural patterns.

## 2. Methods

### 2.1. Participants

Participants were recruited through online campus community boards and the university’s psychology research credit system. Participants included 43 individuals (13 males, 30 females), ranging in age from 19 to 32 years (M = 22.76, SD = 2.95), with the highest proportion (28.6%) being 21 years old.

Exclusion criteria included significant health problems, abnormal visual or auditory perception, and the potential for adverse emotional reactions to specific types of stimuli (e.g., bugs, blood, or ghosts). Prior to participation, all individuals provided informed consent in accordance with institutional ethical guidelines (IRB No. JBNU 2024-09-013-002).

### 2.2. Stimuli

The researchers selected an initial pool of video clips from publicly available YouTube content, focusing on those that clearly conveyed target emotions. A total of 24 clips were chosen to elicit emotional responses, with six assigned to each of four categorical emotional states. Specifically, each category, High-arousal Positive (HP), Low-arousal Positive (LP), High-arousal Negative (HN) and Low-arousal Negative (LN), included six videos.

To ensure that the video clips accurately reflected the intended valence and arousal dimensions, a preliminary validation was conducted prior to the main experiment. A pilot study involving seven participants was carried out to validate and select affective video stimuli. Participants rated candidate clips on a 7-point Likert scale using adjectives corresponding to four predefined affective categories: high-arousal positive, low-arousal positive, high-arousal negative, and low-arousal negative. The emotion scale was constructed from 14 terms systematically varying across valence (positive, neutral, negative) and arousal (high vs. low). Positive emotions included enjoyment, happy, calm, peaceful, relief, and satisfied; the neutral category was represented by surprised; and negative emotions included angry, annoyed, anxious, afraid, disgusted, sad, and melancholy. In terms of arousal, high-arousal emotions comprised enjoyment, happy, surprised, angry, annoyed, anxious, afraid, and disgusted, whereas low-arousal emotions included calm, peaceful, relief, satisfied, sad, and melancholy. For each affective category, six clips with the highest mean ratings were selected. When a clip received similarly high scores across multiple categories, it was assigned to the category in which it obtained the highest mean. The final set of selected clips yielded mean scores ranging from 5.50 to 6.00 for high-arousal positive, 5.75 to 6.54 for low-arousal positive, 3.83 to 4.26 for high-arousal negative, and 3.17 to 4.14 for low-arousal negative. The final selected video clips ranged in duration from 30 seconds to 1 minute.

### 2.3. EEG Recording

EEG data were recorded using Muse 2 headset, a wearable four-channel EEG device. The electrodes were positioned at AF7, AF8 (frontal sites), TP9 and TP10 (temporal sites) according to the international 10-20 system (Jasper, 1958). The reference electrode was placed at FPz.

Raw EEG signals were acquired via the *MindMonitor* app at a sampling rate of 256 Hz. During recording, the app applied basic band filtering, enabling the extraction of standard EEG frequency bands, namely Delta (1-4 Hz), Theta (4-8 Hz), Alpha (8-13 Hz), Beta (13-30 Hz), and Gamma (30-50 Hz)

### 2.4. Procedure

Participants were seated in a quiet room and instructed to minimize movement during the session. After the EEG device was fitted and signal quality was verified, the experiment began immediately.

Participants watched a series of emotional video clips on a laptop screen while wearing the EEG headset. EEG data were recorded continuously throughout the entire viewing period. The video clips were presented in a fixed pseudorandomized order to prevent videos from the same condition from appearing consecutively thereby reducing potential order and carryover effects, this sequence was identical for all participants. After each video clip, participants also provided subjective ratings of their emotional experience using the same emotion scales that were used in the pilot test.

### 2.5. Statistical analyses

Statistical analyses were performed using MATLAB R2024b.

#### 2.5.1. Classifications

To evaluate how accurately dimensions of core affect could be decoded from behavioral and EEG responses, we conducted two-way classification analyses using a support vector machine (SVM) model. Models were trained and tested to discriminate between positive and negative valence, as well as high and low arousal. These analyses were conducted at both within-participant and cross-participant levels to capture distinct aspects of response consistency.

Before conducting EEG classification, the relatively short duration of each segment (30–60 seconds) was taken into consideration. Principal component analysis (PCA) was deemed less suitable in this context due to its sensitivity to noise and the instability of covariance estimation in short time series. Accordingly, EEG features were extracted by averaging the temporal dimension of each trial.

In within-participant classification, the model was trained and tested using data from the same individual. For each participant, the classifier was trained on both behavioral and EEG data from 23 of the 24 emotional video clips (training set) and evaluated on data from the remaining clip (test set). This procedure was repeated 24 times so that each stimulus served once as the test case, constituting a 24-fold cross-validation for each participant.

In cross-participant classification, the model was trained on both behavioral and EEG data from 42 participants (training set) and evaluated on data from the remaining participant (test set). This procedure was repeated 43 times so that each participant served once as the test case, constituting a 43-fold cross-validation. This approach evaluates the generalizability of emotion-related behavioral ratings and EEG patterns across individuals.

For both classification types, we calculated classification accuracy for each participant and assessed whether the average accuracy across participants exceeded chance level. Statistical significance was assessed using p-values derived from the binomial distribution.

#### 2.5.2. Multidimensional Scaling (MDS)

To investigate the representational structure of behavioral and EEG responses to emotional stimuli, we applied MDS with vector fitting. For each participant, a 24 × 24 (stimuli × stimuli) correlation matrix was constructed to capture the pairwise similarity of responses to the 24 video clips. In the behavioral domain, these matrices were derived from affective ratings, averaged across repeated presentations of each stimulus. In the neural domain, EEG features were obtained by averaging across the temporal dimension of each trial, after which pairwise correlations were computed between neural responses to each stimulus pair. Individual matrices from both modalities were then averaged across participants to generate group-level similarity matrices, which were subsequently used as input for MDS.

To evaluate their correspondence with theoretical emotion models, Procrustes rotation was applied to align the empirical spaces with predefined design coordinates reflecting core affect dimensions (Table 2). Subsequently, correlations were computed between the MDS-derived coordinates and the design values to quantify the degree of correspondence among behavioral ratings, EEG-based representations, and the hypothesized emotional structure.

**Table 1.**
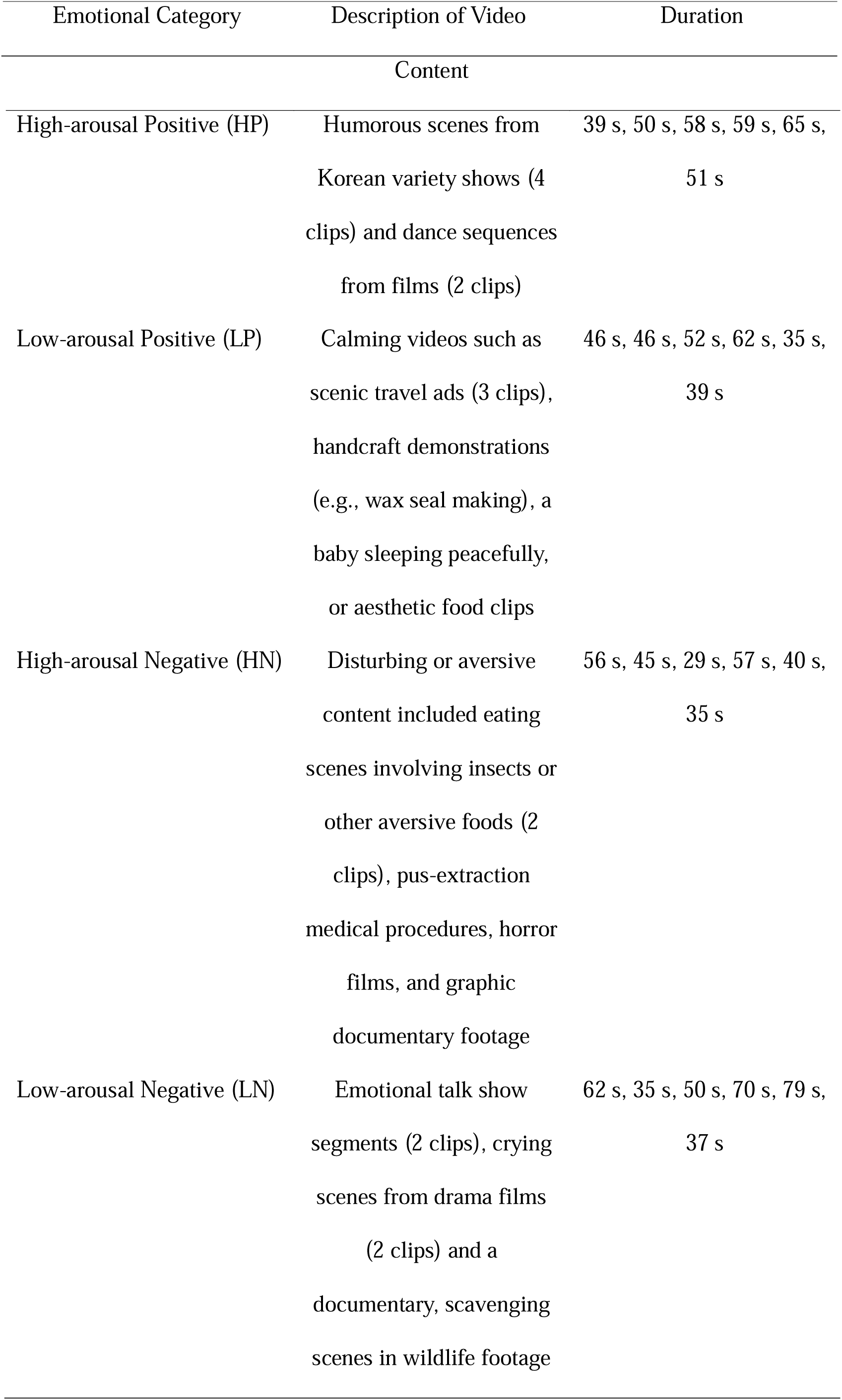
Description and examples of video clips used in each affective stimulus categories.

**Table 2.** Design matrix for affective stimulus categories.

| Affective category | Valence | Arousal |
| --- | --- | --- |
| High-arousal Positive (HP) | 1 | 1 |
| Low-arousal Positive (LP) | 1 | -1 |
| High-arousal Negative (HN) | -1 | 1 |
| Low-arousal Negative (LN) | -1 | -1 |

*2.5.3. ISCs analysis and 3-way repeated-measures ANOVA*

**Table 3.** Results of three-way repeated-measures ANOVA on ISC values.

| Source | df | F | p |
| --- | --- | --- | --- |
| Frequency band | 2.18, 91.61 | 18.89 | < .001 |
| Electrode site | 2.20, 92.21 | 37.06 | < .001 |
| Emotion | 2.63, 110.37 | 2.32 | 0.087 |
| Emotion × Frequency band | 5.84, 245.22 | 5.95 | < .001 |
| Emotion × Electrode site | 5.68, 238.38 | 7.29 | < .001 |
| Frequency band × Electrode site | 4.19, 176.06 | 5.54 | < .001 |
| Emotion × Band ×<br>Electrode | 13.58, 570.21 | 1.97 | 0.019 |

Note: Summary of the three-way repeated-measures ANOVA results on ISC values, with within-subject factors of stimulus type, frequency band, and electrode site. Greenhouse-Geisser corrections were applied to all degrees of freedom. Significant effects ( $p < .05$ ) are highlighted in bold.

To better understand the relationship between the behavioral ratings, EEG responses, and the extracted dimensions, vector fitting was performed on the rotated MDS solution (Borg & Groenen, 2005). Each vector corresponded to either an item from the behavioral rating scales or a channel from the EEG data. Vector length indicated the strength of the association, whereas vector orientation denoted the predicted value corresponding to the orthogonal projection onto the vector. These parameters reflected the relative weighting of each dimension.

Using regression coefficients (expressed as ratios of the two-dimensional coordinates) and coefficients of determination (R²), derived from the regression model line lengths for each dimension, were then used to map the 14 affective responses and 20 EEG features (5 frequency bands × 4 electrode sites) within the core affect space representing the video stimuli.

#### 2.5.3. ISCs analysis and 3-way repeated-measures ANOVA

ISC is a method used to quantify the degree of similarity in neural responses across individuals who are exposed to the same time-locked stimulus, such as a video, narrative, or music (Jang & Kim, 2025; Kim et al., 2023). A high ISC value indicates that participants’ neural activities are responding in a temporally synchronized manner, suggesting shared cognitive or emotional processing of the stimulus (Hasson et al., 2004).

To apply this framework to the current study, we assessed the similarity of participants’ EEG responses to emotional video stimuli using the ISC method. Specifically, we employed a leave-one-out approach (Nastase et al., 2019), wherein each participant’s EEG time series was correlated with the average EEG signal of all other participants, excluding their own data. This procedure allowed us to evaluate how closely each individual’s neural response aligned with the group’s collective response. EEG data for ISC computation consisted of continuous time series covering the entire duration of each emotional video clip. To accommodate minor length mismatches across participants within a category, signals were trimmed to the common minimum length per category prior to correlation. Any missing samples were replaced with zero. ISC was computed separately for each frequency band, electrode site, and stimulus, resulting in one ISC value per participant for each condition. We computed Pearson correlations between the target participant and the leave-one-out group mean separately for each frequency-band × electrode feature (5 bands × 4 electrodes = 20 features). Specifically, we correlated matching feature columns and retained the diagonal of the resulting 20×20 correlation matrix as the feature-wise ISC vector. The computation was repeated iteratively for all participants and across all emotional video conditions.

To capture frequency-specific neural dynamics, ISC values were calculated separately for each frequency band (Delta, Theta, Alpha, Beta, and Gamma) and each electrode site (TP9, AF7, AF8, and TP10). Following ISC computation, Fisher’s r-to-z transformation was applied to the correlation coefficients to improve the normality of the data distribution and to facilitate subsequent parametric analyses, such as ANOVA. After Fisher z-transformation, stimulus-level ISC values were averaged within each emotional category (HP, LP, HN, and LN) to yield one mean ISC per category × frequency band × electrode site for each participant. These participant-level ISC arrays (4 × 5 × 4) were then used as input for the repeated-measures ANOVA.

To examine the effects of experimental conditions on ISC values, we conducted a series of three-way repeated-measured ANOVA. The within-subject factors were emotional categories (HP, LP, HN, and LN), Frequency Band, Electrode Site Greenhouse-Geisser corrections were applied when Mauchly’s test indicated violations of sphericity. A significant threshold of *p < .05* was used for omnibus tests.

To further explore interaction effects involving the Electrode factor, we performed planned contrasts comparing Temporal (TP9 & TP10) vs. Frontal (AF7 & AF8) sites, and Left (TP9 & AF7) vs. Right (TP10 & AF8) hemispheric sites. These contrasts were designed to test whether ISC patterns varied by anatomical location, reflecting functionally relevant spatial differences in EEG synchrony across conditions.

## 3. Results

### 3.1. Behavioral results

#### 3.1.1. Classifications

The mean accuracy of within-participant classification for valence was significant (M = .958, p < .001, Figure 1), indicating that within individuals, behavioral ratings reliably predicted the valence of the video being watched. Similarly, the mean accuracy for classifying arousal was also significant (M = .786, p < .001), suggesting that behavioral ratings also predicted the arousal level of the video. Notably, accuracy for predicting valence was significantly higher, *t*(42) = 12.47, *p* < .001.

**Figure 1.**
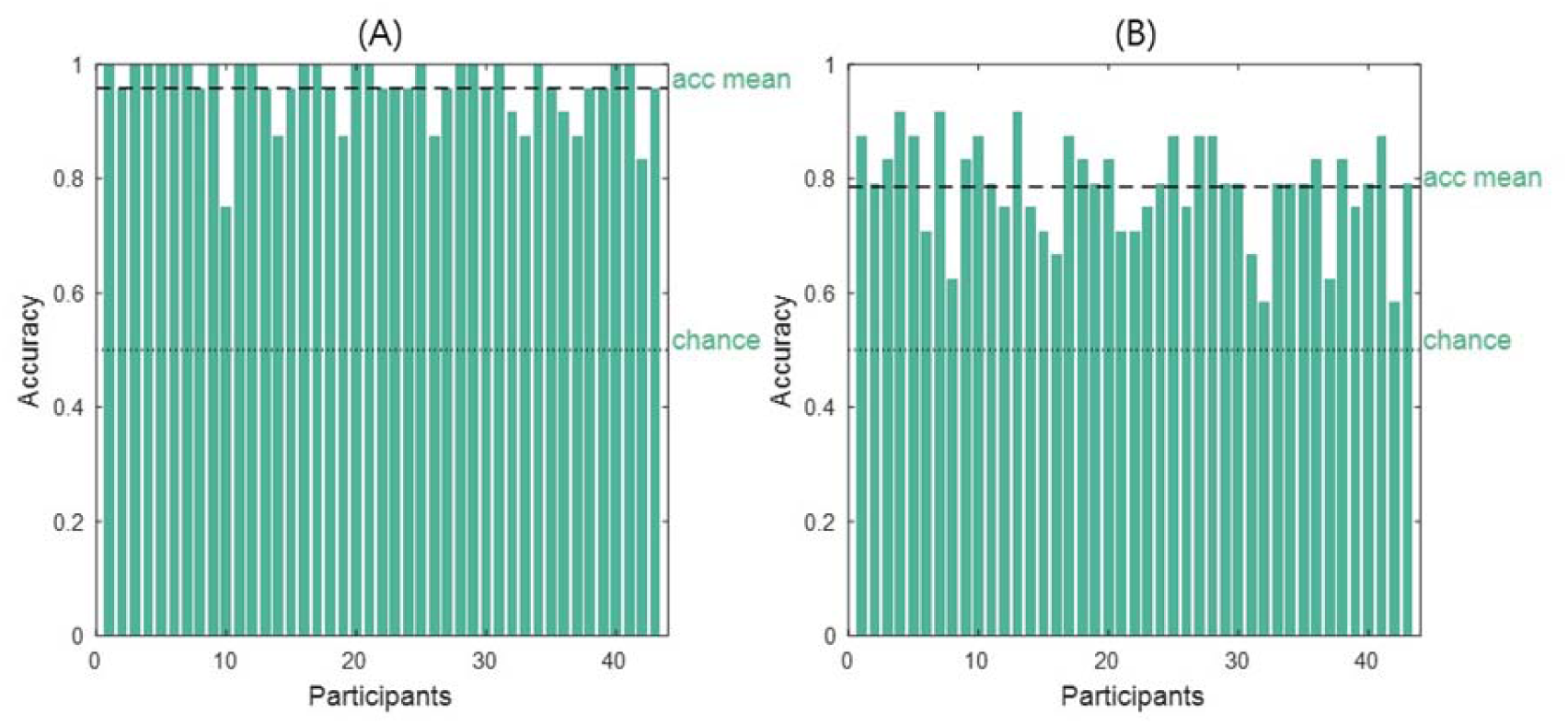
Results of within-participant classification. The dashed line indicates the group mean accuracy, and the dotted line denotes the chance level (0.5). Panel (A) shows the 2-way classification accuracies for valence, and Panel (B) shows the 2-way classification accuracies for arousal. Each bar indicated individual accuracy.

The mean accuracy of cross-participant classification for valence was significant (M = .964, p < .001, Figure 2), indicating that within individuals, behavioral ratings reliably predicted the valence of the video being watched. Similarly, the mean accuracy for classifying arousal was also significant (M = .863, p < .001), suggesting that behavioral ratings also predicted the arousal level of the video. Accuracy for predicting valence was significantly higher, *t*(42) = 8.379, *p* < .001.

**Figure 2.**
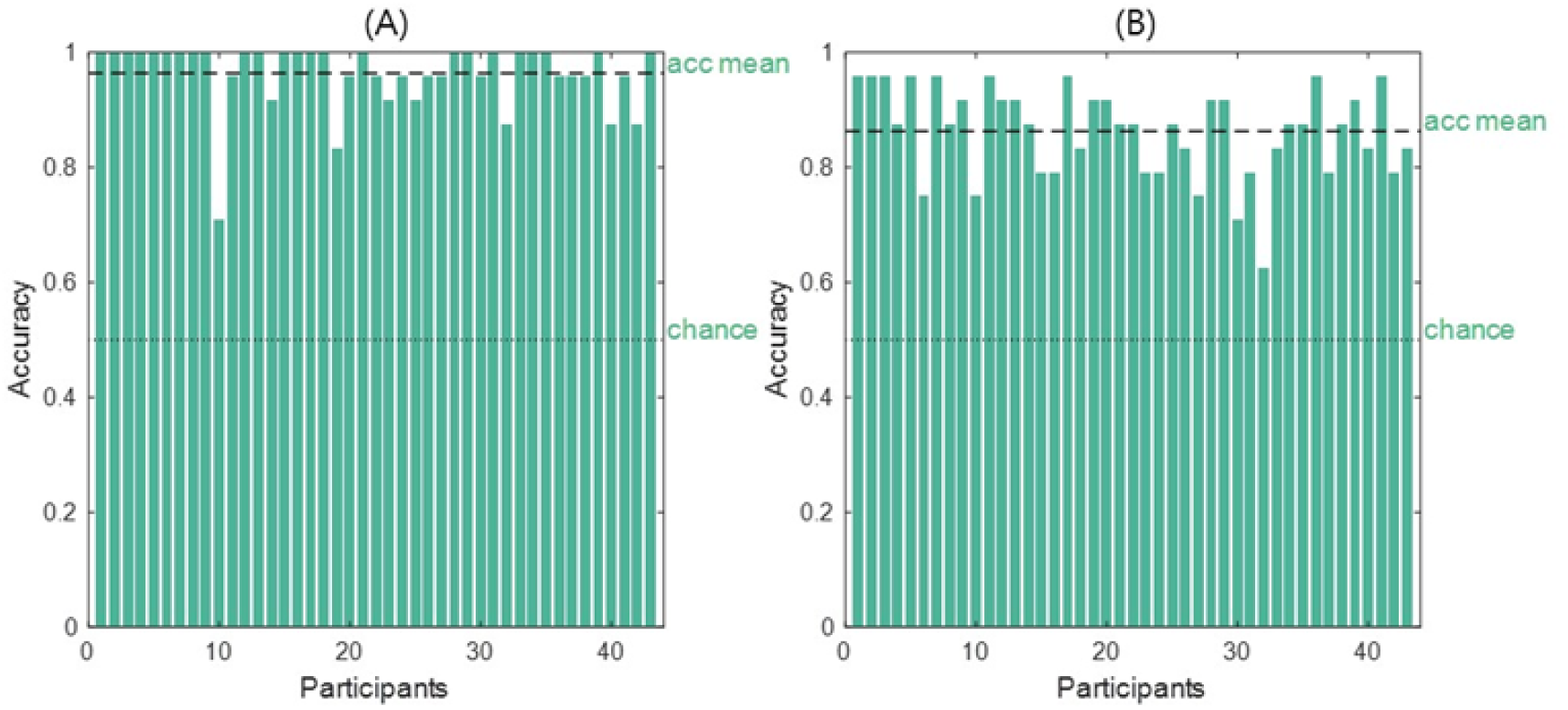
Results of cross-participant classification. The dashed line indicates the group mean accuracy, and the dotted line denotes the chance level (0.5). Each bar indicated individual accuracy.

#### 3.1.2. MDS

MDS was applied to visualize the similarity structure of behavioral ratings across emotional conditions. The two-dimensional MDS solutions were rotated to better align with psychological dimensions (Table. 2). The first dimension showed a a significant correlation with valence (*r* = 0.989, p<.001), while the second dimension also exhibited a significant correlation with arousal (*r* = 0.723, p<.001) (Figure 3).

**Figure 3.**
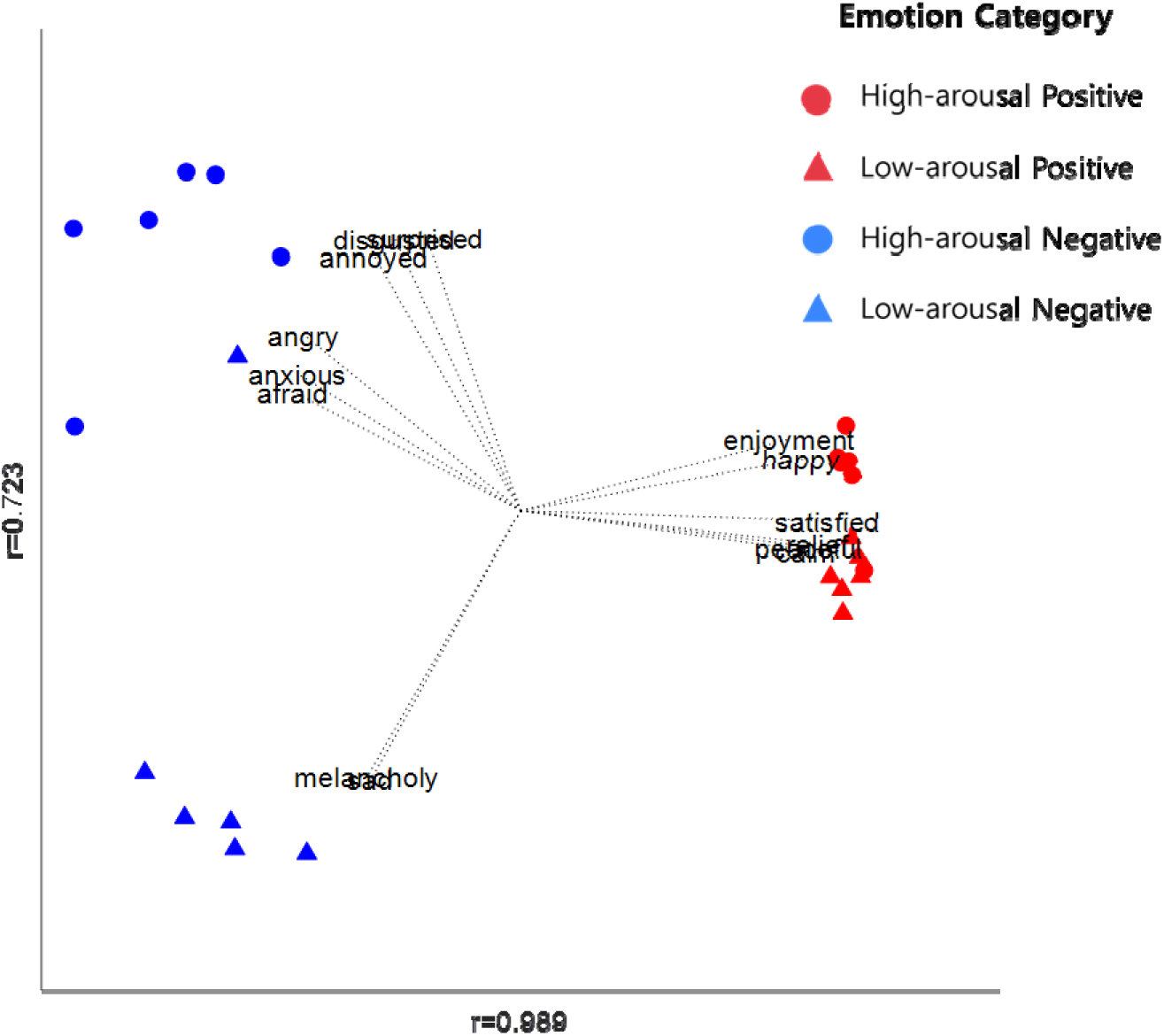
Two-dimensional MDS solution showing the spatial distribution of emotional conditions with vector fitting based on behavioral ratings. Overlaid vectors represent contributions from emotion scales, obtained through vector fitting onto the MDS solution rotated to align with the design matrix. The orientations of these vectors reflect the degree to which each component covaries with the stimulus configuration along the valence and arousal dimensions. Correlation coefficients with the target dimensions are shown, valence (*r* = 0.989), and arousal (*r* = 0.723).

The resulting affective space was consistent with the circumplex model (Russell & Barrett, 1999; Russell, 2003), with emotional conditions distributed primarily along the valence and arousal dimensions. Conditions with opposing valence (e.g., positive vs. negative) appeared linearly separable in the low-dimensional space. This dimensional configuration suggests that affective states can be reliably distinguished based on a low-dimensional signature.

Vector fitting further supported this structure, as individual emotion terms were positioned in accordance with their hypothesized affective categories. For instance, happiness and enjoyment clustered near the positive region of the space, while anger, fear, and disgust were located closer to the negative, high-arousal quadrant. Low-arousal negative emotions, such as melancholy and sadness, were clearly separated toward the lower arousal side. In contrast, low-arousal positive states, such as calm and peaceful, were located adjacent to the positive axis but lower in arousal.

Interestingly, the separation between high- and low-arousal states was more pronounced within the negative domain than within the positive domain. Negative conditions displayed a clear contrast between high-arousal states (e.g., angry, anxious) and low-arousal states (e.g., sad, melancholy). In contrast, positive conditions showed less distinction between high- and low-arousal emotions, with most positive states clustering relatively close to each other. This asymmetry indicates that arousal plays a stronger discriminative role in structuring negative affective space, while valence dominates the organization of positive affective states.

### 3.2. EEG results

#### 3.2.1 Classifications

The mean accuracy of the 2-way within-participant valence classification (Figure 4) was 0.712 (*p* = .011), also significantly above the chance level of 0.50, whereas the mean accuracy of the 2-way arousal classification was 0.583 (*p* = .271), which did not reach significance. These results indicate that, within participants, classification was significant for the valence dimension, whereas classification along the arousal dimension was less robust. Given that the within-participant analysis was based on a limited number of trials (24 in total), caution is warranted in interpreting the null finding for arousal. Accuracy for predicting valence was significantly higher, *t*(42) = 5.133, *p* < .001.

**Figure 4.**
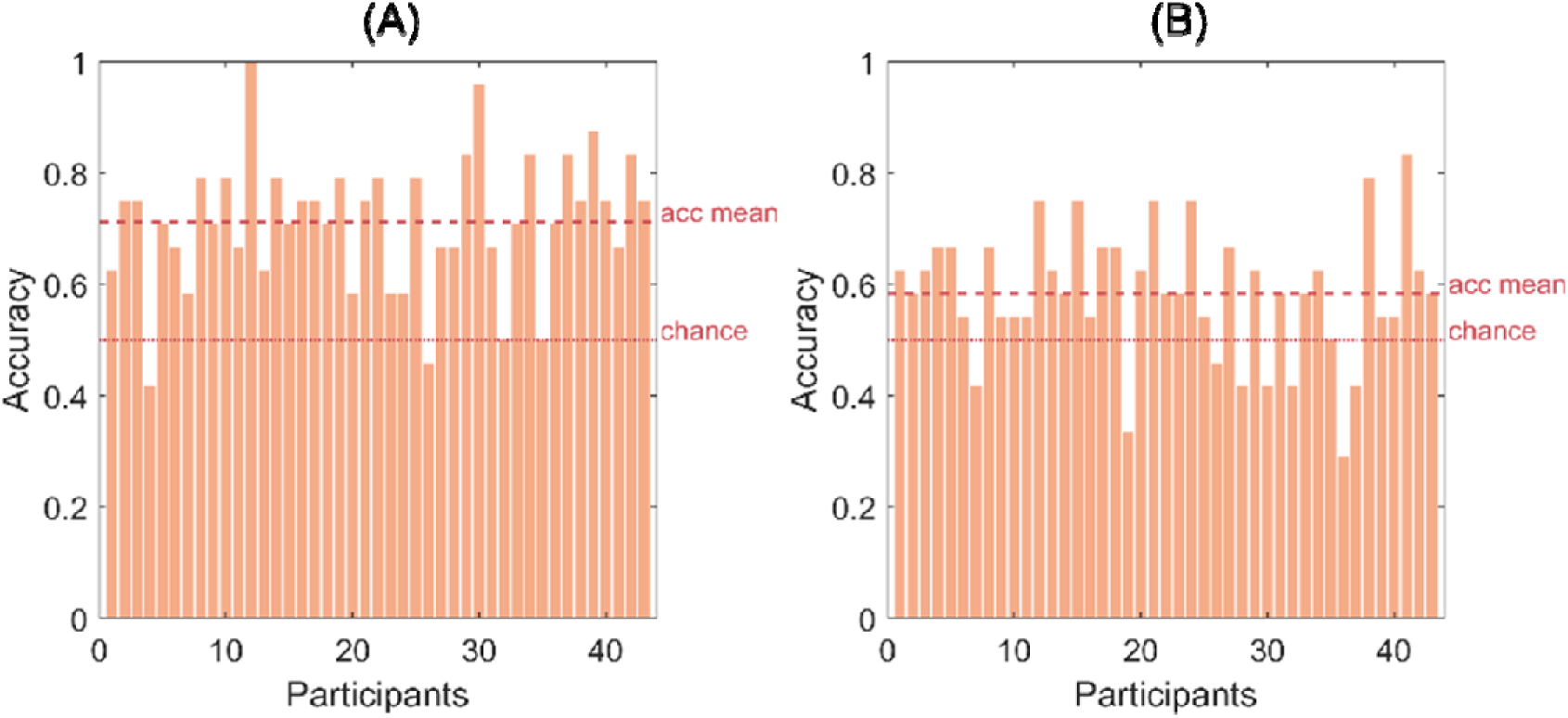
Within participant classification results. Panel A shows the 2-way classification accuracies for valence, with the red dashed line indicating the group mean accuracy and the dotted line denoting the chance level (0.5). Panel B shows the 2-way classification accuracies for arousal. Each bar indicated individual accuracy.

The mean accuracy of the 2-way valence cross-participant classification (Figure 5) was 0.602 (*p* < .001), also significantly above the chance level of 0.50, and the mean accuracy of the 2-way arousal classification was 0.642 *(p* < .001), which likewise exceeded the chance level of 0.50. These results indicate that, across participants, classification was significant for the valence and arousal dimensions. There was no significant difference in accuracy for valence and arousal, *t*(42) = -1.445, *p* = .156.The successful cross-participant classification of valence and arousal categories suggests that affect-related EEG activation patterns are robust and generalizable across individuals. This inter-individual consistency implies that neural representations of affect are shared across participants, as reflected in the overlap of discriminative activation patterns. These findings align with the observed consistency in the spatial distribution of informative EEG features across individuals. Notably, cross-participant classification was performed using the same set of features as within-participant classification, indicating that shared affective representations were robust enough to support generalization across individuals.

**Figure 5.**
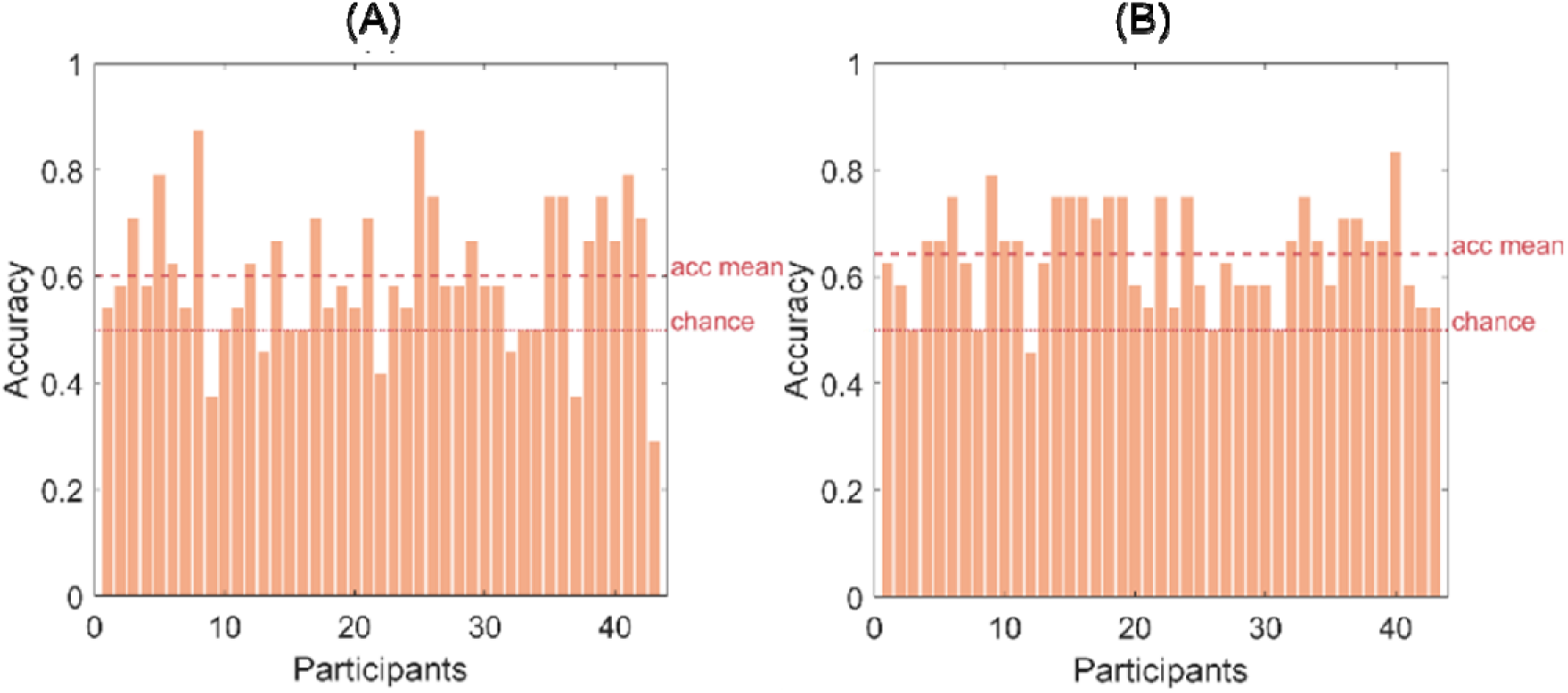
Cross participant classification results. Panel A shows the 2-way classification accuracies for valence, with the red dashed line indicating the group mean accuracy and the dotted line denoting the chance level (0.5). Panel B shows the 2-way classification accuracies for arousal. Each bar indicated individual accuracy.

#### 3.2.2. MDS

MDS was applied to visualize the similarity structure of EEG responses across emotional conditions. The two-dimensional MDS solutions were rotated to better align with psychological dimensions (Table 2). The first dimension was significantly correlated with valence (*r* = 0.806, p<.001), while the second dimension was not significantly correlated with arousal (*r* = 0.312, p=.16), suggesting that the spatial configuration was primarily driven by valence-related neural activity (Figure 6).

**Figure 6.**
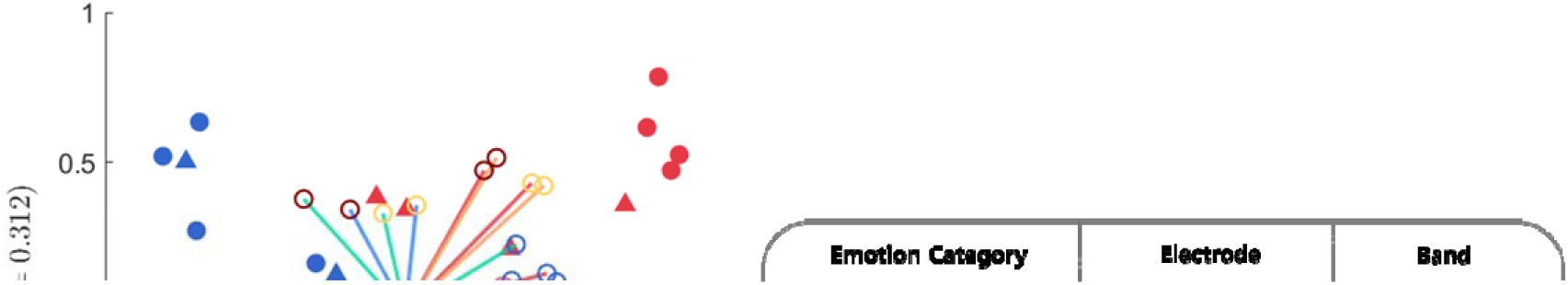
A two-dimensional MDS solution showing the spatial distribution of emotional conditions with vector fitting. Overlaid vectors represent contributions from selected EEG channels (e.g., TP9, TP10) and frequency bands (e.g., Delta, Theta, Alpha), obtained through vector fitting onto the MDS solution rotated to align with the design matrix. The orientations of these vectors reflect the degree to which each component covaries with the stimulus configuration along the valence and arousal dimensions. Correlation coefficients with the target dimensions are shown, valence (*r* = 0.806), and arousal (*r* = 0.312).

The resulting affective space was consistent with the circumplex model (Russell & Barrett, 1999; Russell, 2003), with emotional conditions distributed primarily along the valence dimension and partially along the arousal dimension. This neural representational structure closely mirrored the behavioral results, indicating that both measures capture a common underlying affective organization. However, compared to behavioral data, the EEG-based representation showed a weaker separation along the arousal dimension.

In the sensor space, vectors corresponding to TP9 and TP10 (red and orange lines) were oriented toward the positive end of the valence axis, indicating consistent involvement of temporal regions. In the spectral domain, delta, theta, and alpha bands (red, blue, and green text, respectively) also showed vector orientations toward the positive valence axis. This alignment was supported by vector fitting results (*r* = 0.388), suggesting that these components contributed systematically to the valence dimension.

### 3.4. ISC analysis

ISC values across all frequency bands, electrode sites, and stimuli generally range between 0 and 0.2 (Figure 7). To assess whether mean ISC values differed significantly across conditions, we performed a three-way repeated-measures ANOVA with within-subject factors of stimulus type, frequency band, and electrode site.

**Figure 7.**
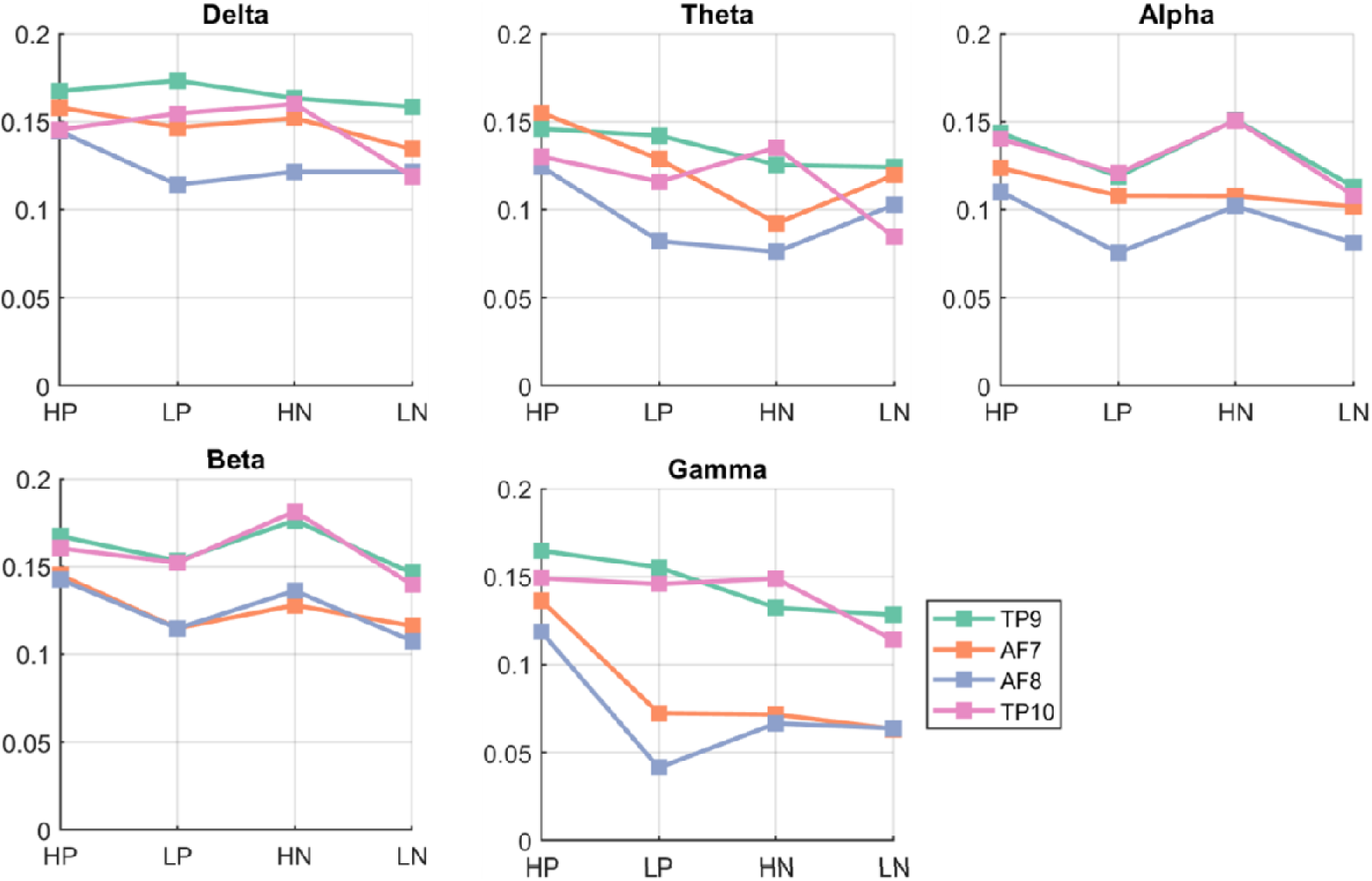
Mean inter-subject correlation (ISC) values across stimulus types (HP, LP, HN, LN), frequency bands (Delta-Gamma), and electrode sites (TP9, AF7, AF8, TP10). Each panel represents a frequency band, and lines indicate ISC values across stimuli for each electrode. While the plots visually suggest interactions among stimulus, band, and electrode, the three-way repeated-measures ANOVA confirmed the statistical significance of these effects.

The results revealed significant main effects of frequency band, *F*(2.18, 91.61) = 18.89, *p* < .001, and electrode site, *F*(2.20, 92.21) = 37.06, *p* < .001, indicating that each independently influenced ISC values. However, the main effect of emotion was not statistically significant, *F*(2.63, 110.37) = 2.32, *p* = .087. Both two-way and three-way interaction effects were significant, Stimulus x Band: *F*(5.84, 245.22) = 5.95, *p* < .001; emotion × electrode: *F*(5.68, 238.38) = 7.29, *p* < .001; band × electrode: *F*(4.19, 176.06) = 5.54, *p* < .001; emotion × band × electrode: *F*(13.58, 570.21) = 1.97, *p* = .019, suggesting that ISC values varied depending on specific combinations of stimulus type, frequency band, and electrode location. All degrees of freedom were corrected using the Greenhouse-Geisser method.

To further examine regional differences in ISC values, *a priori contrasts* were conducted. The left and right hemispheres were compared by averaging across all frequency bands and stimulus types. ISC values were significantly higher in the left hemisphere than in the right, *t* = 7.15, *p* < .001, *M*_left_ = 0.16, *M*_right_ = 0.14. Similarly, a contrast between frontal (AF7, AF8) and temporal (TP9, TP10) electrodes revealed significantly greater ISC values in the temporal regions, *t* = 6.18, *p* < .001, *M*_frontal_ = 0.13, *M*_temporal_ = 0.17, indicating a robust anterior-posterior difference in neural synchrony across participants. These contrasts were defined a priori based on established theories of functional asymmetries between hemispheres and anterior-posterior brain regions.

## 4. Discussion

This study examined behavioral and neural responses to four affective categories of high-arousal positive, low-arousal positive, high-arousal negative, and low-arousal negative using the commercially available EEG device Muse 2, based on the core affect model. Multivariate analyses such as MDS, classification analyses, and ISC were performed to investigate the underlying affective representations.

Behavioral classification analyses indicated that self-reported emotion ratings reliably predicted both the valence and arousal of video stimuli within and across participants, with especially high accuracy for valence. These findings suggest that valence may represent a more stable and predictable dimension of affective experience than arousal. EEG analyses likewise showed that neural signals could reliably predict valence and arousal. While valence classification remained significantly above chance in both within- and cross-participant analyses, arousal classification was less robust, particularly within participants where accuracy did not reach significance. A parsimonious explanation is that the null finding for arousal in the within-participant analysis reflects the limited number of trials, which constrained statistical power. Thus, the result is better interpreted as a methodological limitation due to insufficient data rather than as evidence for the absence of arousal-related neural signatures.

The apparent inconsistency between behavioral and physiological results can be understood in light of the different processes each measure captures. Behavioral reports directly reflect subjective experience, whereas EEG may be more sensitive to very early or fine-grained stages of neural processing (Reid et al., 2024). When participants viewed video stimuli, immersion, attention, and emotional intensity were likely heightened, contributing to clear and consistent behavioral ratings. In contrast, EEG signals are more susceptible to noise and artifacts, potentially obscuring subtle affective patterns. This distinction helps explain why valence was more reliably predicted across both behavioral and neural measures, whereas arousal appeared less stable and exhibited greater inter-individual variability (Kuppens et al., 2013).

Next, MDS was employed to further explore the representational structure of affective responses. The MDS results for both behavioral and EEG data demonstrated clear affective separation along the valence axis, suggesting that valence may be represented in a more stable and spatially consistent behavioral and neural format. These results contrast with findings from Yang et al. (2024), in which arousal was more clearly represented. This discrepancy may stem from differences in stimulus structure and experimental design that helped compensate for the limitations of time-averaged EEG analysis. In the prior study, stimuli were relatively long in duration (2–3 minutes) and narrative in nature, designed to elicit clear and sustained arousal responses. As a result, participants’ physiological arousal may have been more consistently maintained throughout the stimulus period, allowing for clearer distinctions between arousal levels even when using the same EEG feature extraction method. These findings suggest that effective representation of arousal in EEG data depends not only on analytical techniques but also on the temporal structure and elicitation characteristics of the stimuli. The interaction between these factors may play a critical role in successful arousal decoding.

Recent studies similarly report that valence is more stably represented than arousal in both behavioral and physiological/EEG data. For example, Jessup et al. (2025) used MDS to map the structure of emotion items and found clearer separation along the valence dimension than along arousal. Similarly, Bruin et al. (2024), comparing self-reports and physiological signals across stressors, found that valence ratings were more consistent across measures, whereas arousal showed greater variability. Garg et al. (2022) likewise demonstrated with portable EEG that valence can be decoded with higher reliability than arousal under similar conditions.Consistent with these findings, the EEG results for arousal condition showed relatively weak separability in both the MDS and 2-way within-participant classification results. This may be attributed to the interaction between the analytical characteristics of the EEG and the structure of the stimuli. In the present study, EEG features were extracted using a time-averaged approach, without incorporating temporal information. Arousal is often dynamic in nature, and its expression can vary significantly over time. Static, mean-based features may fail to capture brief but intense neural responses associated with arousal. While traditional EEG analyses often rely on static, mean-based features such as average band power, these measures may obscure critical temporal information embedded in the signal. Affective arousal, in particular, is characterized by brief but intense neural responses that emerge within the first few hundred milliseconds following stimulus onset. Olofsson et al. (2008) demonstrated that high-arousing stimuli elicit rapid ERP components, such as early posterior negativity (EPN), N2, P300, and late positive potential (LPP), that evolve dynamically across time. Because these responses are transient and time-locked to stimulus presentation, static power averages risk attenuating or missing them entirely. This underscores the importance of adopting time-resolved approaches that can preserve the evolving temporal structure of emotion-related brain activity. Notably, Zheng and Lu (2015) demonstrated that features reflecting time-frequency dynamics of EEG signals achieved higher accuracy in emotion classification tasks. In this context, the use of static, averaged features in the present study may have limited the ability to capture transient arousal signals, thereby constraining classification performance based on arousal.

ISC was introduced in the analysis to quantify the degree of similarity in neural responses across individuals exposed to the same time-locked stimulus. The results revealed greater neural synchronization in the left frontal region compared to the right. However, the present study yielded generally low ISC values, with most averages falling below 0.2. This can be interpreted as follows. The importance of temporal dynamics is not only critical for capturing arousal-related EEG features but also extends to ISC analysis. In fact, the generally low ISC values observed in this study may reflect not just the challenges of detecting arousal, but also limitations in the temporal structure of the affective stimuli. ISC is particularly sensitive to the temporal structure of neural responses, which in turn requires stimuli with sufficient duration to allow for meaningful cross-participant synchronization. When stimuli are too brief, they may elicit only transient, weakly synchronized responses, or in some cases, no measurable synchronization at all. This notion is supported by previous studies such as Hasson et al. (2004) and Dmochowski et al. (2012), which reported high and stable ISC values using stimuli with longer durations ranging from 6 to 30 minutes. These findings suggest that longer stimuli enable the alignment of temporally extended cognitive and affective processes across individuals, thereby increasing ISC sensitivity. In contrast, the short video clips (30-60 seconds) used in the present study may not have provided sufficient temporal depth for stable cross-brain synchronization to emerge. Thus, stimulus duration likely acted as a limiting factor in the current ISC analysis.

Despite these methodological and conceptual limitations, the present study offers several important contributions to the affective neuroscience field, particularly in feasibility, scalability, and ecological applicability. First, it contributes to the growing body of literature evaluating the applicability of the core affect model using consumer-grade EEG devices. By demonstrating that valence-based differentiation of emotional stimuli is robust even with a low-cost, wearable EEG system, this study supports the feasibility of theory-driven emotion research outside laboratory settings. Second, the study provides a quantitative analysis of how affective stimuli influence inter-subject neural synchronizations, as indexed by ISC, offering new insight into the consistency of emotional processing across individuals. Importantly, this study highlights the possibility of conducting theoretically grounded research in ecologically valid settings without relying on high-end laboratory equipment. As such, it offers a foundation for future studies aiming to develop low-cost, portable EEG-based methods for emotion monitoring in real-world environments.

Taken together, these findings suggest that commercial EEG devices, when paired with appropriate analytical strategies, can serve as practical tools for investigating affective processes across individuals. This work not only advances our understanding of affective representation and neural synchrony but also lays the groundwork for scalable, accessible, and context-sensitive emotion research in everyday settings.

